# Distinct evolutionary dynamics of horizontal gene transfer in drug resistant and virulent clones of *Klebsiella pneumoniae*

**DOI:** 10.1101/414235

**Authors:** Kelly L Wyres, Ryan R Wick, Louise M Judd, Roni Froumine, Alex Tokolyi, Claire L Gorrie, Margaret M C Lam, Sebastián Duchêne, Adam Jenney, Kathryn E Holt

## Abstract

*Klebsiella pneumoniae* (*Kp*) has emerged as an important cause of two distinct public health threats: multidrug resistant (MDR) healthcare-associated infections^1^ and community-acquired invasive infections, particularly pyogenic liver abscess^2^. The majority of MDR hospital outbreaks are caused by a subset of *Kp* clones with a high prevalence of acquired antimicrobial resistance (AMR) genes, while the majority of community-acquired invasive infections are caused by ‘hypervirulent’ clones that rarely harbour acquired AMR genes but have high prevalence of key virulence loci^3–5^. Worryingly, the last few years have seen increasing reports of convergence of MDR and the key virulence genes within individual *Kp* strains^6^, but it is not yet clear whether these represent a transient phenomenon or a significant ongoing threat. Here we perform comparative genomic analyses for 28 distinct *Kp* clones, including 6 hypervirulent and 8 MDR, to better understand their evolutionary histories and the risks of convergence. We show that MDR clones are highly diverse with frequent chromosomal recombination and gene content variability that far exceeds that of the hypervirulent clones. Consequently, we predict a much greater risk of virulence gene acquisition by MDR *Kp* clones than of resistance gene acquisition by hypervirulent clones.

---

*Kp* MDR evolution is largely driven through acquisition of AMR genes on diverse mobilisable plasmids^7^ which are particularly prevalent among the subset of clones that have become globally disseminated and frequently cause hospital outbreaks^8^; e.g. clonal group (CG) 258 which is implicated in global spread of the *Kp* carbapenemases^9^. *Kp* pathogenicity is driven by a wide array of interacting factors^10 –12^ that are present in all strains, including the type III fimbriae (*mrk*) and the surface polysaccharides (capsule and lipopolysaccharide (LPS))^12, 13^ which exhibit antigenic variation between strains. The majority of hypervirulent *Kp*, distinguished clinically as causing invasive infections even outside the hospital setting^14^, are associated with just two^4, 15^ of the >130 predicted capsular serotypes^16^, K1 and K2, that are considered particularly antiphagocytic and serum resistant^15, 17^. Hypervirulent *Kp* are also associated with high prevalence of several other key virulence factors; the *rmpA*/*rmpA2* genes that upregulate capsule expression to generate hypermucoidy^18, 19^; the colibactin genotoxin that induces eukaryotic cell death and promotes invasion to the blood from the intestines^20, 21^; and the yersiniabactin, aerobactin and salmochelin siderophores that promote survival in the blood by enhancing iron sequestration^11,22–24^.

Yersiniabactin synthesis is encoded by the *ybt* locus, which is usually mobilised by an integrative, conjugative element known as ICE*Kp.* It is present in ∼40% of the general *Kp* population and seems to be frequently acquired and lost from MDR clones^25^. Fourteen distinct *ybt*+ICE*Kp* variants are recognised, one of which also carries the colibactin synthesis locus (*clb*)^25^. In contrast, the salmochelin (*iro*), aerobactin (*iuc*) and *rmpA*/*rmpA2* loci are usually co-located on a virulence plasmid^26, 27^. These loci are much less common in the *Kp* population (<10% prevalence each) and until recently were rarely reported among MDR strains^3^^3,5^.

The reasons for the apparent separation of MDR and hypervirulence are unclear but there are growing reports of convergence from both directions, i.e. hypervirulent strains gaining MDR plasmids^28–33^ and MDR strains gaining a virulence plasmid plus/minus an ICE*Kp*^6,30,34^. Most such reports are sporadic, but in 2017 Gu and colleagues described a fatal outbreak of MDR, carbapenem-resistant *Kp* belonging to CG258 that had acquired ICE*Kp* in the chromosome plus *iuc* and *rmpA2* on a virulence plasmid^6^. The report fuelled growing fears of an impending public health disaster in which highly virulent MDR strains may be able to spread in the community, causing dangerous infections that are extremely difficult to treat^35^. However, there remain significant knowledge gaps about *Kp* evolution that limit our ability to understand the severity of this public health threat, and to evaluate the relative risks of convergence events.

Here we report a comparative analysis of genome evolutionary dynamics between MDR and hypervirulent clones, leveraging a curated collection of 2265 *Kp* genomes to identify 28 common clones with at least 10 genomes in each (total 1092 genomes in 28 clones, 10–266 genomes per clone, see **Supplementary Tables 1,2** and Figure 1a,b). First we explored the distribution of key virulence loci and resistance determinants for 10 distinct antimicrobial classes across the total population (**Methods**). The presence of the virulence plasmid (with or without ICE*Kp*) was negatively associated with the presence of acquired AMR genes: the majority (n=77/88) of genomes harbouring the virulence plasmid contained 0–1 acquired AMR genes, while the distributions were much broader for genomes without the virulence plasmid plus/minus ICE*Kp* (median 1, interquartile range (IQR) 0-9, p < 1×10^−10^ for both pairwise Wilcoxon Rank Sum tests). Interestingly, among the genomes without the virulence plasmid the distribution of acquired AMR genes was slightly shifted towards higher numbers in genomes harbouring ICE*Kp* (median 1 vs 1, IQR 0-8 vs 0-10, p < 1×10^−8^; see **Supplementary Figure 1**). Next we calculated the proportion of genomes harbouring the virulence and AMR loci within each of the 28 common clones (Figure 1c). Hierarchical clustering of the virulence locus data defined a group of 6 hypervirulent clones including the previously described CG23, CG86 and CG65^3^, each of which harboured the virulence plasmid-associated genes *iuc, iro* and/or *rmpA/rmpA2* at high frequency (31– 100%). Hierarchical clustering of the AMR data defined a group of 8 MDR clones, including the outbreak associated CG258, CG15 and CG147^8^, each with a high frequency (≥56%) of genomes encoding acquired resistance determinants for ≥3 drug classes (in addition to ampicillin to which all *Kp* share intrinsic resistance via the chromosomally encoded SHV-1 beta-lactamase). As expected, AMR genes were rare among the hypervirulent clones, with the exception of CG25 in which 11 of 16 genomes harboured ≥4 acquired AMR genes. The *iuc, iro* and *rmpA*/*rmpA2* loci were rare (<12% frequency) among the MDR clones and those not assigned to either group (‘unassigned’ clones); however, the *ybt* locus was frequently identified across the spectrum of clones as has been reported previously^25^.

**Figure 1:**
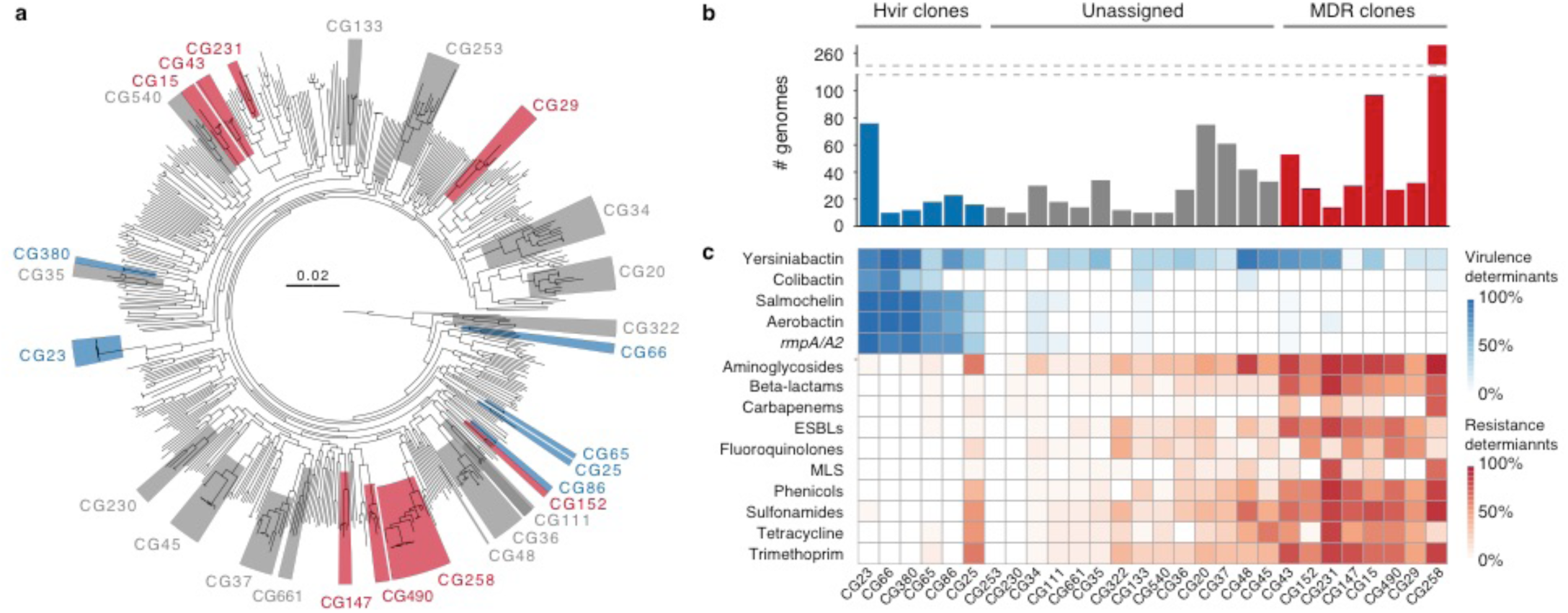
*K. pneumoniae* clones investigated in this study. **a)** Phylogenetic tree inferred using maximum likelihood for *K. pneumoniae* genomes selected from our curated collection to represent the 509 distinct 7-gene chromosomal multi-locus sequence types. Phylogenetic clusters (monophyletic groups) were defined using patristic distance (cut-off = 0.04). Clusters corresponding to clones included in comparative analyses are marked; blue, hypervirulent; grey, unassigned; red, multi-drug resistant. **b)** Total number of genomes included in comparative analyses, coloured by clone type as above. Note that sample sizes exceed the number of isolates shown in the tree for the corresponding clones. **c)** Distribution of virulence and resistance determinants by clone. Intensity of box shading indicates the proportion of genomes harbouring the key virulence loci (blue) or acquired genes conferring resistance to different classes of antimicrobials (red), as per inset legends. Hypervirulent (Hvir) clones were identified by hierarchical clustering of virulence locus data. Multi-drug resistant (MDR) clones were identified by hierarchical clustering of resistance data. AMR, antimicrobial resistance; *rmpA*/*A2*, regulators of mucoid phenotype; ESBLs, extended spectrum beta-lactams; MLS, macrolide, lincosamide and streptogramin B antibiotics.

We used Gubbins^36^ to identify putative chromosomal recombination imports within each clone and calculated r/m (the ratio of single nucleotide variants introduced by homologous recombination relative to those introduced by substitution mutations), which ranged from 0.02–25.50 (Figure 2a, **Supplementary Table 1**). With the exception of CG25, the hypervirulent clones generally exhibited lower r/m values (median 1.15), while the MDR clones trended towards higher values (median 5.47) although the differences were not statistically significant (Kruskall-Wallis test p = 0.07). Recombination events were not evenly distributed across chromosomes: in 19/28 clones ≥50% of the chromosome was not subject to any recombination events, while the maximum recombination load in each clone ranged from mean 1.1–47.6 events (Figure 2b, **Supplementary Figure 2**).

**Figure 2:**
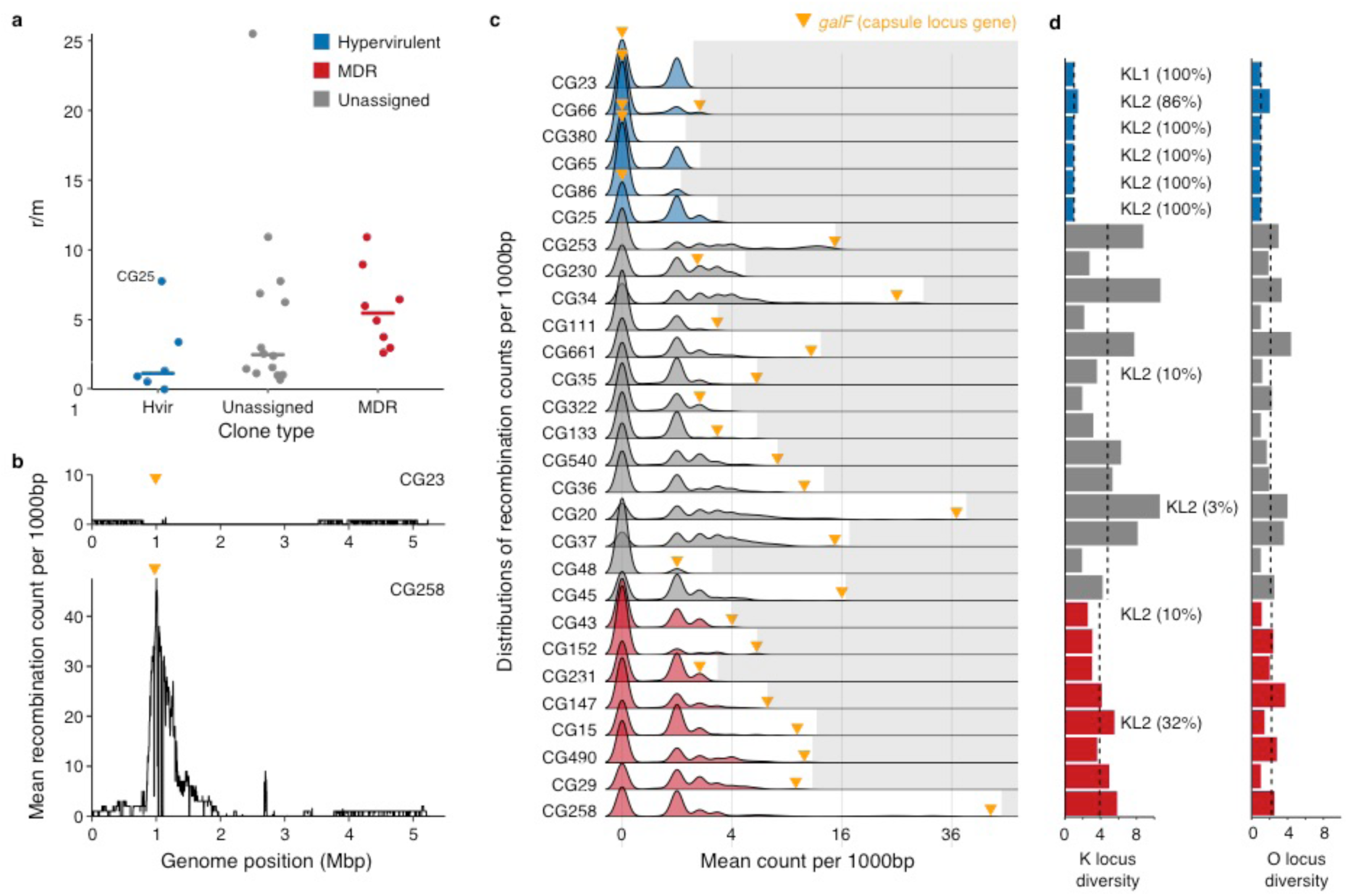
Recombination dynamics, capsule (K) and LPS antigen (O) locus diversity. **a)** Scatter plot showing the ratio of single nucleotide polymorphisms introduced by recombination vs mutation (r/m) for each clone grouped by clone type (n = 6, 14 and 8 for the hypervirulent, unassigned and MDR groups, respectively). Bars indicate median values. **b)** Example plots showing mean recombination counts per base calculated over non-overlapping 1000 bp windows of the chromosome for hypervirulent CG23 and multidrug resistant CG258. The latter has a distinct peak in recombination counts around the K/O loci (marked by the yellow arrow). **c)** Density plots showing the distributions of mean recombination counts per base calculated as in **(b)**. For each row, grey shading marks values outside the distribution of that clone and the yellow arrow indicates the value for the window containing *galF*, the 5’-most K locus gene. Plots are coloured by clone type as above. **d)** K and O locus diversities by clone (effective Shannon’s diversities). Clones harbouring KL1 or KL2 encoding the highly serum resistant capsule types K1 and K2, respectively are marked (numbers in parentheses indicate the percentage of successfully typed genomes harbouring the locus). Bars are coloured by clone type as above. Dashed lines indicate median values for each clone type.

In many cases there was a major peak defining a recombination hot-spot at the capsule (K) and adjacent LPS antigen (O) biosynthesis loci (see e.g. CG258 in Figure 2b, and **Supplementary Figure 2**). Among the 17 clones with ≥1 detectable recombination hotspot (arbitrarily defined as mean recombination count of ≥5 per base calculated over non-overlapping 1000 bp windows), the *galF* K locus gene was ranked among the top 2% recombination counts in 16 clones (Figure 2c and **Supplementary Figure 2**). Consistent with these findings, 20 clones were associated with ≥3 distinct K loci and 11 clones were also associated with ≥3 O loci (Figure 2d and **Supplementary Figures 3**, 4). Together these data indicate that the capsule and LPS are subject to strong diversifying selection. However, this was not the case for the hypervirulent clones, which were associated with low K and O locus diversity: five out of the six had just one K and one O locus type (either KL1 or KL2, plus O1/O2v1 or O1/O2v2) and showed no evidence of recombination events affecting *galF* (Figure 2c,d **Supplementary Figures 3, 4**).

**Figure 3:**
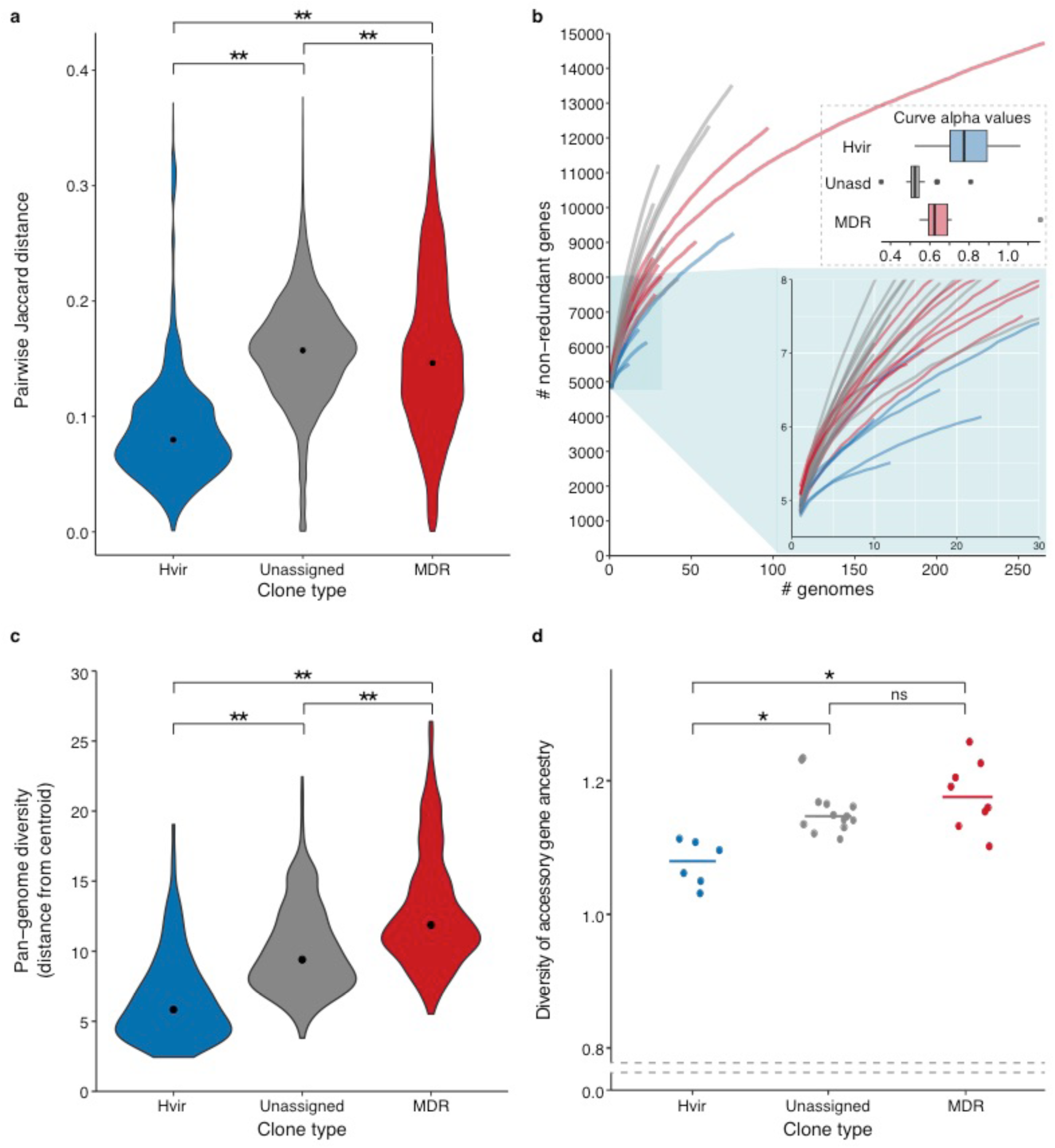
Gene content diversity. **a)** Pairwise gene content Jaccard distances were calculated for all pairs of genomes within each clone and are summarised by clone type (n = 7150, 15754, 86228 pairs for the hypervirulent, unassigned and MDR groups, respectively). Black points indicate median values. **b)** Gene accumulation curves were generated independently for each clone using the rarefy function in the R Vegan^12^ package to analyse each gene content matrix, and are coloured by clone type. The upper inset box shows the distributions of alpha values calculated using the R micropan^13^ package, where alpha <1 indicates an open pan-genome and alpha >1 indicates a closed pan-genome. The lower inset box shows a magnified view for up to 30 genomes. **c)** Violin plots showing the distributions of Euclidean distances from clone centroids for each genome, calculated from the gene content matrix after decomposition to 40 dimensions (n = 157, 390 and 547 for the hypervirulent, unassigned and MDR groups, respectively). Black points indicate median values. **d)** Scatter plot showing ancestral diversity of accessory genes for each clone grouped by clone type. Accessory genes were identified as those present in <95% genomes. Each gene was assigned to a putative ancestral origin using Kraken v0.10.6, genus level assignments were used to calculate Shannon’s diversity indices (n = 6, 14 and 8 for the hypervirulent, unassigned and MDR groups, respectively). Horizontal lines indicate median values. Note that the y-axis is broken. For all panels, brackets indicate Wilcoxon Rank Sum tests of pairwise group comparisons; ns, not significant; *, p < 0.01; **, p < 1×10^−15^.

**Figure 4:**
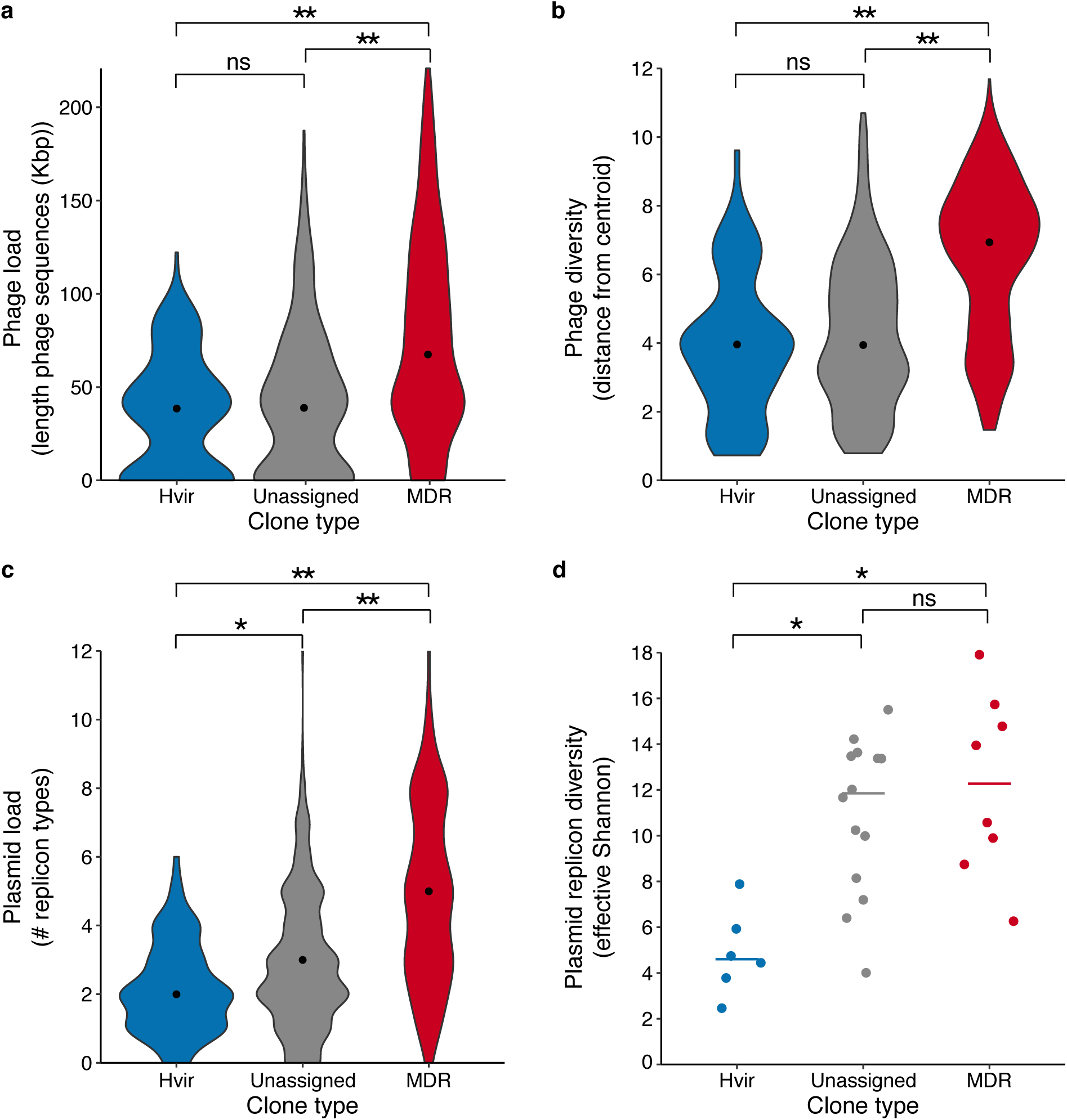
Phage and plasmid diversity. **a)** Violin plots showing the distributions of the total length (kbp) of phage sequence identified per genome. **b)** Violin plots showing the distributions of Euclidean distance to clone centroids calculated from the phage gene content matrix decomposed into 25 dimensions. **c)** Violin plots showing the distributions of plasmid replicon count per genome (note that perfectly co-occurring replicons are counted once only). **d)** Effective Shannon’s diversities for plasmid replicons, by clone. Horizontal lines indicate median values (n = 6, 14 and 8 for the hypervirulent, unassigned and MDR groups, respectively). For all violin plots, data points represent individual genomes (n = 157, 390 and 547 for the hypervirulent, unassigned and MDR groups, respectively) and black points indicate median values. For all panels, brackets indicate Wilcoxon Rank Sum tests of pairwise group comparisons; ns, not significant; *, p < 0.01; **, p < 1×10^−15^.

The key selective drivers for K/O locus diversity are not known, but the mammalian immune system is unlikely to play a major role since *Kp* live ubiquitously in the environment and are opportunistic rather than obligate human pathogens^37^. Instead, phage and/or protist predation are likely candidates^38^. Numerous capsule specific *Kp* phage have been reported^39, 40^ and ecological modelling supports a key role for phage-induced selective pressures in maintaining surface polysaccharide diversity in free-living bacteria^41^. The relative lack of diversity among the hypervirulent clones may suggest that they are not subject to the same selective pressures, perhaps indicating some sort of ecological segregation. This possibility is intriguing and could explain the separation of hypervirulence and MDR, by limiting opportunities for horizontal gene transfer between MDR and hypervirulent clones. Isolates representing both clone types have been identified among diverse host-associated niches^31–42,43^ but it is not possible to determine any particular ecological preference due to the lack of systematic sampling efforts to-date. An alternative explanation is that the hypervirulent clones are subject to some sort of mechanistic limitation for chromosomal recombination, that in turn limits surface polysaccharide diversity and the acquisition of other chromosomally encoded accessory genes, as have recently shown to be frequently acquired by CG258 strains^44^. If so, we may also expect a general trend towards lower gene content diversity in the hypervirulent clones.

To assess overall gene content diversity we conducted a pan-genome analysis using Roary^45^. Jaccard gene content distances were generally lower for genome pairs within hypervirulent clones than the MDR or unassigned clones, suggesting the former have less diverse pan-genomes (p < 1×10^−15^ for each pairwise Wilcoxon Rank Sum test, Figure 3a). Supporting this trend, the hypervirulent clones were associated with comparatively shallow pan-genome accumulation curves (Figure 3b). In order to quantify the differences in these curves we fitted the pan-genome model proposed by Tettelin and colleagues^46^, and derived an alpha value for each clone (**Supplementary Table 1**), whereby values <1 indicate an open pan-genome and >1 indicate a closed pan-genome. Consistent with previous data showing extensive gene content diversity within the *Kp* species^5^, all but two clones had alpha values below 1. The exceptions were hypervirulent CG380 (alpha = 1.06) and MDR CG231 (alpha = 1.16). There was a general trend towards higher alpha (i.e. less open) among the hypervirulent clones (median alpha = 0.77, IQR 0.70–0.89), although the difference was only statistically significant in comparison to the unassigned clones (median alpha = 0.52, IQR 0.51–0.54, p = 0.005) and not the MDR clones (median alpha = 0.62, IQR 0.59–0.69, p = 0.23, Figure 3b).

It is well known that large groups of accessory genes can be linked on the same mobile element (e.g. large conjugative MDR plasmids that are common in the MDR clones, or the virulence plasmids characteristic of the hypervirulent clones), so a single gain or loss event may have a large effect on gene-based measures such as pairwise Jaccard distances and accumulation curves. Hence we used a principal component analysis (PCA) to generate a metric that is less sensitive to the correlation structure in the gene content data (see **Methods**). The PCA transformed the accessory gene content matrix comprising 1092 genomes vs 39375 genes into coordinates in a 40-dimensional space. These 40 axes captured >60% of the variation in accessory gene content and were used to calculate the Euclidean distance of each genome to its clone centroid. The resulting distributions of distances provided further support that the MDR and unassigned clones display greater gene content variation than hypervirulent clones (Figure 3c; p < 1×10^−15^, 2 d.f., Kruskal-Wallis test; p < 1×10^−15^ for each pairwise Wilcoxon Rank Sum test) and suggest this is associated with a greater frequency of horizontal gene transfer events rather than a similar number of events introducing larger changes in gene content. In addition, the putative ancestry of accessory genes (see **Methods**) was more diverse among MDR and unassigned clones than the hypervirulent clones, supporting that the latter are subject to a more limited range of partners for horizontal gene transfer (Wilcoxon Rank Sum tests: hypervirulent vs MDR, p = 0.0027; hypervirulent vs unassigned, p = 1×10^−4^; MDR vs unassigned, p = 0.38; Figure 3d, **Supplementary Table 1**).

To further explore differences in common sources of accessory gene diversity, we assessed phage and plasmid diversity. For each genome we summed the length of genomic regions identified as phage by VirSorter^47^ (range 0–221 kbp, **Supplementary Table 2**) and used a PCA of phage-associated gene content to calculate distance to clone centroids as for the total pan-genome (Figure 4a,b and **Supplementary Figure 6**). Hypervirulent clones showed similar phage load and diversity to the unassigned clones (Figure 4a,b; Wilcoxon Rank Sum tests, hypervirulent vs unassigned: load, p = 0.15; diversity, p = 0.15), whereas MDR clones were generally associated with higher load and diversity than both the hypervirulent and unassigned groups (p < 1×10^−15^ for each pairwise comparison). Although these analyses were dependent on the quality and breadth of the underlying viral sequence database which may be subject to species bias, we have no reason to expect that this would be skewed with respect to MDR over hypervirulent clones. Hence it is clear that *Kp,* and in particular the MDR clones, are subject to frequent attack by diverse phage.

Unfortunately it is not possible to reliably identify plasmid sequences from draft genome assemblies^48^. Instead we used plasmid replicon and relaxase (*mob*) typing as indicators of plasmid load and diversity. Each genome contained 0–12 of 69 uniquely distributed replicon markers and 0–23 *mob*-positive assembly contigs (detected by screening against the PlasmidFinder^49^ database and *mob* PSI-BLAST^50, 51^, respectively; **Supplementary Table 2**). MDR and unassigned genomes harboured a greater number of replicon markers than hypervirulent genomes, largely driven by low replicon loads in CG23 that was overrepresented among the hypervirulent genomes (Figure 4c,d, **Supplementary Figure 7**; Wilcoxon Rank Sum tests: hypervirulent vs unassigned, p = 1.5×10^−4^; MDR vs unassigned, p < 1×10^−15^; MDR vs hypervirulent, p < 1×10^−15^). There were no significant differences between the hypervirulent and unassigned groups for counts of *mob-*positive contigs per genome, and comparatively small differences between the MDR and unassigned or hypervirulent groups (Wilcoxon Rank Sum test: MDR vs unassigned, p = 1×10^−6^; MDR vs hypervirulent, p <1×10^−6^, **Supplementary Figure 7**).

Comparison of effective Shannon’s diversity of replicon profiles indicated that the hypervirulent clones harbour less plasmid diversity than either of the unassigned or MDR clones (not driven solely by CG23, see Figure 4d, **Supplementary Figure 7,** Wilcoxon Rank Sum tests: hypervirulent vs MDR, p = 0.0013; hypervirulent vs unassigned, p = 0.0015; MDR vs unassigned, p = 0.44). Similar trends were seen for effective Shannon’s diversity of *mob* types but the differences were not statistically significant after Bonferroni correction for multiple testing (n=3 tests), a finding that is not surprising given that far fewer *mob* types have been defined and that only ∼48% completely sequenced Enterobacteriaceae plasmids deposited in GenBank could be *mob* typed (whereas ∼83% could be replicon typed)^50^.

While these data are subject to the biases of the underlying databases within which clinically relevant (MDR and virulence) plasmids are overrepresented, they are also consistent with the findings above regarding overall pan-genome diversity. These data imply that MDR clones frequently acquire and lose plasmids, consistent with the high plasmid diversity reported previously for ST258^52, 53^ and several others^7^, and with data from recent investigations of *Kp* circulating in hospitals which showed that individual plasmids transferred frequently between clones^54, 55^. In contrast, the hypervirulent clones were associated with comparatively low plasmid diversity, mirrored by generally narrow plasmid load distributions. Taken together these data imply that hypervirulent clones acquire novel plasmids infrequently but can stably maintain them. For example, we recently estimated that the virulence plasmid, which by definition is highly prevalent in these clones, has been maintained for >100 years in CG23^31^. In addition, laboratory passage experiments have shown that hypervirulent strains can maintain MDR plasmids introduced *in vitro*^29^, and we showed that a horse-associated subclade of CG23 has maintained a single MDR plasmid for at least 20 years^31^.

The combination of infrequent plasmid acquisitions and limited chromosomal recombination suggests that hypervirulent clones may be subject to particular constraints on DNA uptake and/or integration. One possible explanation is that these *Kp* clones possess enhanced defences against incoming DNA such as CRISPR/Cas or restriction-modification (R-M) systems. However, our genome data reveals no significant differences in either system (see **Supplementary Text**, **Supplementary Figures 8**–**11**). Alternatively, the key virulence determinants themselves, or other proteins encoded on the virulence plasmid, may play a role. Two variants of the virulence plasmid predominate among hypervirulent clones and share limited homology aside from the *iuc, iro* and *rmpA* loci^27^. It seems unlikely that a siderophore system would influence DNA uptake, however it is conceivable that upregulation of capsule expression by *rmpA*^18, 56^ may play a role by exacerbating the inhibitory effect of the capsule.

Capsule expression has been associated with a comparative reduction in *Kp* transformation frequency *in vitro*^57^ and in a natural *Streptococcus pneumoniae* population^58^. Additionally, the capsule is known to conceal the LPS^59^ which, together with the OmpA porins, are considered key target sites for attachment of conjugative pili during the initial phases of mate-pair formation^60, 61^. Hence we speculate that overexpression of the capsule in hypervirulent clones may result in a reduction of DNA uptake. Given that capsule types differ substantially in their thickness and polysaccharide composition^18, 62^, it is also likely that their influence on DNA uptake is type dependent. The K2 capsule, which is associated with five of the 6 hypervirulent clones investigated here is considered among the thicker capsule types^18^ and thus may have a comparatively greater influence. We used our genome data to test this hypothesis by comparing the genomic diversity of KL2 and non-KL2 genomes within MDR CG15, the only clone with sufficient KL2 and non-KL2 genomes for comparison. CG15-KL2 genomes formed a deep branching monophyletic subclade consistent with long-term maintenance of KL2 for an estimated 34 years (see **Supplementary Text and Supplementary Figures 12 and 13**). This KL2 subclade showed a comparatively low rate of recombination (r/m 0.58 vs 6.75) and more limited gene content diversity than the rest of the clone (p = 0.0004, **Supplementary Text and Supplementary Figure 12**). However, gene-content diversity in the KL2 subclade of CG15 was higher than that of the hypervirulent clones (**Supplementary Figure 12**), perhaps due to the absence of the *rmpA* capsule upregulator. Thus the genome data support our hypothesis and should motivate future laboratory studies of this phenomenon; a task that will not be trivial given the low efficiency of *in vitro* transformation for wild-type *Kp* strains^63^, the challenge of identifying suitable selective markers for distinguishing MDR strains, and the sensitivity of conjugation efficiencies to laboratory growth conditions^60^. If confirmed, this would imply that hypervirulent clones are evolutionarily constrained by a key determinant of the hypervirulent phenotype, and as such are self-limited in their ability to adapt to antimicrobial pressure.

Regardless of the mechanisms, our data clearly show that hypervirulent *Kp* clones are less diverse than their MDR counterparts, and suggest that the rate of virulence plasmid acquisition by MDR clones will far exceed the rate of MDR plasmid acquisition by hypervirulent clones. This is particularly worrying from a hospital infection control perspective since many of the MDR clones investigated here appear well adapted to transmission and colonisation in the human population, and are frequent causes of hospital outbreaks^7, 8^. Given the mounting evidence that MDR clones can carry multiple plasmids at limited fitness cost^64–66^ and frequently exchange plasmids with other bacteria^54, 55^, it seems these MDR clones may also be the perfect hosts for consolidation and onwards dissemination of MDR and virulence determinants. The greatest concern is that these determinants will be consolidated onto a single mobile genetic element; indeed mosaic *Kp* plasmids carrying AMR genes plus *iuc* and *rmpA2* have already been reported in an MDR *Kp* clone^30^, and *Escherichia coli* plasmids bearing *iuc, rmpA* and AMR genes have been detected in *Kp*^27^. Whether these convergent strains and plasmids are fit and disseminating is not known. Recent experience with convergent carbapenem-resistant CG258 in China – which retrospective surveillance studies showed was already widely disseminated at the time of the outbreak report^6, 67^ – highlights the ease with which deadly strains can circulate unnoticed. As reports of convergent *Kp* strains continue to increase, the need for global genomic surveillance encompassing clone, AMR and virulence locus information^68^ is clearly greater than ever.

## Methods

### Genome collection and clone definition

We collected and curated 2265 *Kp* genomes, comprising 647 genomes sequenced and published previously by our group^5, 15,16,69^ plus 1623 publicly available genomes^53, 70–76^ as described previously^16^. Genomes were assigned to chromosomal multi-locus sequence types (MLST, as below), and a single representative of each sequence type (ST, n=509) was selected for initial phylogenetics to define clones for further analysis. Sequence reads were mapped to the NTUH-K2044 reference chromosome (accession: NC_012731) using Bowtie v2^77^ and single nucleotide variants were identified with SAMtools v1.3.1^78^ as implemented in the RedDog pipeline (https://github.com/katholt/RedDog). Where genomes were available only as *de novo* assemblies, sequence reads were simulated using SAMtools wgsim^78^ (n=852 genomes, for each of which 2 million x 100bp PE reads were simulated without errors). Allele calls were filtered to exclude sites that did not meet the following quality criteria: unambiguous consensus base calls, phred quality ≥30, depth ≥5 reads but <2-fold mean read depth, no evidence of strand bias. Subsequently, we generated a variable site alignment by concatenating nucleotides at core genome positions, i.e. at positions for which ≥95% genomes contained a base-call with phred quality ≥20. The resulting alignment of 192,433 variable sites was used to infer a maximum likelihood phylogeny with FastTree v2.1.9^79^ (gamma distribution of rate heterogeneity among sites, Figure 1a). Genomes were clustered into 259 phylogenetic lineages (clones) using patristic distance (distance threshold = 0.04). This identified 29 clones (clonal groups, CGs) that were each represented by ≥10 isolates from at least three different countries. One of these (CG82) was subsequently excluded because it uniquely included only historical isolate genomes (dated 1932–1949 or unknown). The remaining 28 clones (totalling 1092 genomes) were subjected to comparative analysis in this study. We refer to each as CGX, where X is the predominant ST in the clone, as per the convention for *Kp*.

For each clone of interest, reads were mapped to a completed chromosomal reference genome belonging to that clone (see **Supplementary Table 1** and below), and variant calling and phylogenetic inference was performed as above. Phylogenies were manually inspected alongside genome source information to identify and de-duplicate clusters of closely related genomes from the same patient and/or known hospital outbreaks. Additional random sub-sampling was applied to CG258, which was otherwise drastically overrepresented in the collection (>700 genomes subsampled to 266 genomes). The final set of clones and genomes used for analyses are listed in **Supplementary Tables 1 and 2**, respectively.

Note that for the initial investigations of virulence and AMR determinant distributions in the broader *Kp* population (shown in **Supplementary Figure 1**) we considered an independent subset of the original curated genome collection (n=1124). This subset was described previously^16^ and was considered more representative of the population diversity because known outbreaks and overrepresented sequence types were subsampled.

### Clonal reference genome selection

Reference genomes for each clone were identified among publicly available completed *Kp* chromosome sequences for each ST represented in the clones of interest. Where there was no suitable publicly available reference genome we selected a representative isolate from our collection, for which Illumina data were available, and generated additional long read sequence data for completion of the genome through hybrid genome assembly (details below). The exception was CG380 for which no suitable reference was publicly available and for which we did not have access to any isolates in our collection. As such, we generated a pseudo-chromosomal reference by scaffolding the *de novo* assembly contigs for genome SRR2098675^76^ (the CG380 genome with the lowest number of contigs) to the most closely related completed genome in our initial phylogeny (NCTC9136, available via the NCTC3000 genomes project website: http://www.sanger.ac.uk/resources/downloads/bacteria/nctc/). Contigs were scaffolded using Abacas^80^ and manually inspected using ACT^81^ (contig coverage ≥20%).

### Long read sequencing and hybrid genome assembly

Novel completed reference genomes were generated for 10 clones (CG25, CG29, CG36, CG253, CG43, CG45, CG152, CG230, CG231, CG661) for which Illumina data were available^5, 82^. Novel long read data were generated on the Pacific Biosciences platform for two isolates (CG231 strain MSB1_8A, and CG29 strain INF206), and an Oxford Nanopore Technologies MinION device for the remaining isolates as described previously^83^. Long read sequence data were combined with the existing short read Illumina data to generate complete hybrid assemblies with Unicycler^84^. The final completed assemblies were deposited in GenBank (accessions listed in **Supplementary Table 1)** and are available in Figshare (see below).

### MLST, virulence and resistance gene screening

Chromosomal MLST, AMR and virulence genes were detected with SRST2^85^ (or Kleborate, available at https://github.com/katholt/Kleborate, for typing assemblies when no sequence reads were available). Sequence reads were assembled *de novo* using SPAdes v3^86^, and Kaptive v0.5.1^16, 87^ was used to determine K and O locus types from assemblies. K and O locus diversities were calculated using the R package Vegan v2.4.3^88^. The indices were converted to effective values to enable direct comparison between clones using the formula described previously^89^; effective Shannon diversity = exp(Shannon diversity).

### Recombination detection

Recombination analysis was performed independently for each clone: single nucleotide variants were identified by mapping and variant calling against the clonal reference genome as above, and a pseudo-chromosomal alignment was used as input for Gubbins v2.0.0^36^, with the weighted Robinson-Foulds convergence method and RAxML^90^ phylogeny inference. The Gubbins output files were used to calculate r/m and mean recombination counts per base, calculated over non-overlapping 1000 bp windows (relative to each clone-specific reference chromosome).

### Pan-genome, plasmid and phage analyses

The SPAdes derived genome assemblies were annotated with Prokka v1.11^91^ and subjected to a pan-genome analysis with Roary v3.6.0^45^ (BLASTp identity ≥95%, no splitting of ‘paralogs’). The resulting gene content matrix comprised 1092 genomes vs 39375 genes (after excluding 1070 core genes present in ≥95% genomes) was used to calculate pairwise Jaccard distances and as input for PCA with the Adegenet R package v2.0.1^92^. Coordinates for the top 40 principal components (PC) were extracted, capturing 61.1% of the variation in the data. We calculated the Euclidean distance from each genome to its clone centroid (the vector of mean coordinate values for that clone), and compared the distributions of distances across clones. Pan-genome accumulation curves were visualised using the R package Vegan v2.4.3^88^ and alpha values were calculated using R package micropan v1.1.2^93^.

Accessory genes were identified as those present in <95% of all 1092 genomes belonging to the 28 clones analysed. For each genome assembly within a clone, the accessory gene sequences were extracted and concatenated into a single multi-fasta file (one per clone) that was used as input for ancestral assignment by Kraken v0.10.6, run with the miniKraken database^94^. For each clone the proportions of accessory genes assigned to distinct genera were used to calculate Shannon’s diversity indices using the R package Vegan v2.4.3^88^.

Phage were identified from genome assemblies using VirSorter v1.0.3^47^, with the highest confidence threshold. The resulting output includes a set of putative phage sequences in fasta format, in which we identified open reading frames (ORFs) using Prokka v1.11^91^. The resulting ORF sequences were clustered into non-redundant phage gene sequences using CD-HIT-EST v4.6.1^95^ (identity ≥95%). BLASTn was used to tabulate the presence/absence of each of the resulting phage genes within the putative phage sequences identified by VirSorter in each genome (identity ≥95%, coverage ≥95%). The resulting gene content matrix was used as input for PCA and centroid distance calculations as described above for all accessory genes (25 PCs were used, capturing 60.0% of the variation in the data).

Plasmid replicons defined in the PlasmidFinder database^49^ were identified from read data using SRST2^85^ and from assemblies using BLASTn (identity ≥80%, coverage ≥50% for both methods). Identically distributed replicons were collapsed into a single entry to minimise the influence of multi-replicon plasmids. Plasmid *mob* types were identified by PSI-BLAST as previously described^50, 51^. Effective Shannon’s diversities were calculated for each clone based on the replicon and *mob* presence/absence matrices, using the R package Vegan v2.4.3^88^ as described above.

### CRISPR/Cas and restriction-modification systems

CRISPR arrays were identified from genome assemblies using the CRISPR Recognition Tool v1.2^96^ and genomes with >3 putative arrays were investigated manually to check for spurious identifications and/or identifications of single arrays split over multiple assembly contigs. Nucleotide sequences for the previously described *Kp cas* genes^97^ were extracted from the NTUH-K2044 reference chromosome (accession: NC_012731.1). Genomes were screened for novel *cas* genes by HMM domain search using the domain profiles developed by Burstein and colleagues^98^ (HMMER v3.1b2, bit score ≥200^99^). A representative set of putative novel *cas* genes was extracted from the genome of isolate INF256 (read accession: ERR1008719, genome assembly available in Figshare) and tBLASTx was used to detect the presence of NTUH-K2044- and INF256-like *cas* genes among all genomes (identity ≥85%, coverage ≥25%). Note that the INF256 *cas* genes were subsequently found to be highly similar to those of strain Kp52.145 reported during the course of this study^100^.

Putative restriction enzymes (REases) were identified from genome assemblies by HMM domain search using the domain profiles developed previously^101, 102^, parameters as above. CD-HIT v4.6.1^95^ was used to cluster the predicted amino acid sequences of these REases such that distinct clusters represented enzymes that are thought to recognise distinct methylated nucleotide motifs^101^; i.e. using amino acid identity thresholds of 80% identity for type I and type III REases; 55% identity for type II REases. To date no suitable threshold has been determined for type IV REases and thus in order to include type IV enzymes in our analyses we used the more conservative 55% identity threshold. Type IIC REase sequences cannot be aligned^101^ and were therefore excluded from the analysis. Nucleotide sequences for a single representative of each REase cluster were used to search all assemblies by BLASTn (identity ≥80%, coverage ≥90%). Only the single best hit was recorded for each region of each genome. In order to assess the DNA recipient potential of each genome, we also used BLASTn to screen a broader sample of the *Kp* population, comprising the 1124 representative non-redundant genomes from our curated collection as described above and previously^16^, plus a further 598 diverse *Kp* genomes published during the course of this project^103 – 106^. A putative donor-recipient pairing was considered compatible if the complete set of REases in the recipient genome were also present in the donor genome (we assume that genomes positive for an REase also carry the corresponding methyltransferase).

### Investigating the impact of the K2 capsule on CG15 recombination and pan-genome diversity

K loci were overlaid onto the CG15 recombination-free maximum likelihood phylogeny, revealing that the KL2 locus was restricted to one of two major subclades (shaded blue and grey in **Supplementary Figure 12**). Recombination dynamics and pan-genome diversity were investigated separately for each of these subclades, using the methods described above. We used BEAST2^107^ to estimate the time to most recent common ancestor (tMRCA) of the KL2 subclade, using as input the recombination-free single nucleotide variant alignment generated by Gubbins (2967 bp). The final analysis included 21 genomes (those for which years of collection were not known were excluded, see **Supplementary Table 2**) and was completed as described previously^31^. Temporal structure was confirmed by date-randomisation tests, which showed that the evolutionary rate derived from the true data did not overlap those derived from any of 20 independent randomisations (**Supplementary Figure 13**).

### Data availability

Individual accession numbers and genotyping data for all genomes included in comparative analyses are listed in **Supplementary Table 2**. Reference genome accessions are listed in **Supplementary Table 1.** Reference genomes, study genome assemblies and annotations, pseudogenome alignments (Gubbins inputs), Gubbins per branch statistics and recombination predictions output files, python script to calculate mean recombination events per base, pan-genome gene content matrix, python code for Euclidean distance calculation, accessory gene ancestor matrix, representative phage gene sequences, phage gene content matrix, representative REase gene nucleotide sequences and CG15-KL2 BEAST xml input files are available in Figshare (https://doi.org/10.26188/5b8cb880dcffc).

## Acknowledgements

This work was supported by a Viertel Foundation of Australia Senior Medical Research Fellowship to KEH, the Bill and Melinda Gates Foundation, Seattle (OPP1175797), and the University of Melbourne.

## Author contributions

LMJ, CLG, MMCL and AJ provided isolates, performed DNA extractions and/or generated sequence data. KLW, RRW, RM, AT and SD analysed the data. KLW and KEH designed the study and wrote the manuscript. All authors contributed to data interpretation, read and commented on the manuscript.

## References

1. World Health Organization. Global priority list of antibiotic-resistant bacteria to guide research, discovery, and devlopment of new antibiotics. (2017).

2. Shon, A. S., Bajwa, R. P. S. & Russo, T. A. Hypervirulent (hypermucoviscous) *Klebsiella pneumoniae*: a new and dangerous breed. Virulence 4, 107–118 (2013).

3. Bialek-Davenet, S. et al. Genomic definition of hypervirulent and multidrug-resistant *Klebsiella pneumoniae* clonal groups. Emerg Infect Dis. 20, 1812–1820 (2014).

4. Brisse, S. et al. Virulent clones of *Klebsiella pneumoniae*: Identification and evolutionary scenario based on genomic and phenotypic characterization. PLoS One 4, e4982 (2009).

5. Holt, K. E. et al. Genomic analysis of diversity, population structure, virulence, and antimicrobial resistance in *Klebsiella pneumoniae*, an urgent threat to public health. Proc Natl Acad Sci U S A. 112, E3574–81 (2015).

6. Gu, D. et al. A fatal outbreak of ST11 carbapenem-resistant hypervirulent *Klebsiella pneumoniae* in a Chinese hospital: a molecular epidemiological study. Lancet Infect Dis. 3099, 1–10 (2017).

7. Navon-Venezia, S., Kondratyeva, K. & Carattoli, A. *Klebsiella pneumoniae*: A major worldwide source and shuttle for antibiotic resistance. FEMS Microbiol Rev. 41, 252–275 (2017).

8. Wyres, K. L. & Holt, K. E. *Klebsiella pneumoniae* population genomics and antimicrobial-resistant clones. Trends Microbiol. 24, 944–956 (2016).

9. Chen, L. et al. Carbapenemase-producing *Klebsiella pneumoniae*: Molecular and genetic decoding. Trends Microbiol. 22, 686–696 (2014).

10. Martin, R. M. & Bachman, M. A. Colonization, infection, and the accessory genome of *Klebsiella pneumoniae*. Front. Cell. Infect. Microbiol. 8, 1–15 (2018).

11. Holden, V., Breen, P., Houle, S., Dozois, C. & Bachman, M. A. Klebsiella pneumoniae siderophores induce inflammation, bacterial dissemination, and HIF-1α stabilization during pneumonia. 7, 1–10 (2016).

12. March, C. et al. Role of bacterial surface structures on the interaction of *Klebsiella pneumoniae* with phagocytes. PLoS One 8, 1–16 (2013).

13. Cortés, G. et al. Molecular analysis of the contribution of the capsular polysaccharide and the lipopolysaccharide O side chain to the virulence of *Klebsiella pneumoniae* in a murine model of pneumonia. Infect Immun. 70, 2583–2590 (2002).

14. Sellick, J. A. & Russo, T. A. Getting hypervirulent *Klebsiella pneumoniae* on the radar screen. Curr Op Microbiol. (2018). doi:10.1097/QCO.0000000000000464

15. Lee, I. R. et al. Differential host susceptibility and bacterial virulence factors driving *Klebsiella* liver abscess in an ethnically diverse population. Sci Rep. 13, 29316 (2016).

16. Wyres, K. L. et al. Identification of *Klebsiella* capsule synthesis loci from whole genome data. Microb. Genomics. 2, (2016).

17. Kabha, K. et al. Relationships among capsular structure, phagocytosis, and mouse virulence in *Klebsiella pneumoniae*. Infect Immun. 63, 847–852 (1995).

18. Wacharotayankun, R. et al. Enhancement of extracapsular polysaccharide synthesis in *Klebsiella pneumoniae* by RmpA2, which shows homology to NtrC and FixJ. Infect Immun. 61, 3164–3174 (1993).

19. Hsu, C. R., Lin, T. L., Chen, Y. C., Chou, H. C. & Wang, J. T. The role of *Klebsiella pneumoniae rmpA* in capsular polysaccharide synthesis and virulence revisited. Microbiology 157, 3446–3457 (2011).

20. Lu, M.-C. et al. Colibactin contributes to the hypervirulence of *pks*+ K1 CC23 *Klebsiella pneumoniae* in mouse meningitis infections. Front. Cell. Infect. Microbiol. 7, 1–14 (2017).

21. Lai, Y. C. et al. Genotoxic *Klebsiella pneumoniae* in Taiwan. PLoS One 9, e96292 (2014).

22. Bachman, M. A. et al. *Klebsiella pneumoniae* yersiniabactin promotes respiratory tract infection through evasion of lipocalin 2. Infect. Immun. 79, 3309–3316 (2011).

23. Hsieh, P., Lin, T., Lee, C., Tsai, S. & Wang, J. Serum-induced iron-acquisition systems and TonB contribute to virulence in *Klebsiella pneumoniae* causing primary pyogenic liver abscess. J Infect Dis. 197, 1717–1727 (2008).

24. Russo, T. A. et al. Aerobactin mediates virulence and accounts for increased siderophore production under iron-limiting conditions by hypervirulent (hypermucoviscous) *Klebsiella pneumoniae*. Infect. Immun. 82, 2356–2367 (2014).

25. Lam, M. M. C. et al. Genetic diversity, mobilisation and spread of the yersiniabactin-encoding mobile element ICEKp in *Klebsiella pneumoniae* populations. MGen. Epub ahead of print. (2018). doi:http://dx.doi.org/10.1101/098178

26. Ramirez, M. S., Traglia, G. M., Lin, D. L., Tran, T. & Tolmasky, M. E. Plasmid-mediated antibiotic resistance and virulence in gram-negatives: the *Klebsiella pneumoniae* paradigm. Microbiol Spectr. 2, 1–15 (2014).

27. Lam, M. C. C. et al. Tracking key virulence loci encoding aerobactin and salmochelin siderophore synthesis in *Klebsiella pneumoniae*. bioRxiv (2018).

28. Surgers, L., Boyd, A., Girard, P. M., Arlet, G. & Decré, D. ESBL-producing strain of hypervirulent *Klebsiella pneumoniae* K2, France. Emerg. Infect. Dis. 22, 1687–1688 (2016).

29. Xie, Y. et al. Emergence of the third-generation cephalosporin-resistant hypervirulent *Klebsiella pneumoniae* due to the acquisition of a self-transferable blaDHA-1-carrying plasmid by an ST23 strain. Virulence 9, 838–844 (2018).

30. Turton, J. F. et al. Virulence genes in isolates of *Klebsiella pneumoniae* from the UK during 2016, including among carbapenemase gene-positive hypervirulent K1-ST23 and ‘non-hypervirulent’ types ST147, ST15 and ST383. J Med Microbiol. 67, 118–128 (2017).

31. Lam, M. M. C. et al. Population genomics of hypervirulent *Klebsiella pneumoniae* clonal group 23 reveals early emergence and rapid global dissemination. Nat Commun. 9, 2703 (2018).

32. Yao, B. et al. Clinical and molecular characteristics of multi-clone carbapenem-resistant hypervirulent (hypermucoviscous) *Klebsiella pneumoniae* isolates in a tertiary hospital in Beijing, China. Int J Infect Dis. 37, 107–112 (2015).

33. Gu, D.-X. et al. Detection of colistin resistance gene *mcr-1* in hypervirulent *Klebsiella pneumoniae* and *Escherichia coli* isolates from an infant with diarrhea in China. Antimicrob Agents Chemother. 60, 5099–5100 (2016).

34. Heinz, E. et al. Emergence of carbapenem, beta-lactamase inhibitor and cefoxitin resistant lineages from a background of ESBL-producing *Klebsiella pneumoniae* and *K. quasipneumoniae* highlights different evolutionary mechanisms. bioRxiv (2018). doi:10.1101/283291

35. Chen, L. & Kreiswirth, B. N. Convergence of carbapenem-resistance and hypervirulence in *Klebsiella pneumoniae*. Lancet Infect Dis. 18, 9–10 (2018).

36. Croucher, N. J. et al. Rapid phylogenetic analysis of large samples of recombinant bacterial whole genome sequences using Gubbins. Nucleic Acids Res. 43, e15 (2015).

37. Podschun, R. & Ullmann, U. *Klebsiella* spp. as nosocomial pathogens: epidemiology, taxonomy, typing methods, and pathogenicity factors. Clin Microbiol Rev. 11, 589–603 (1998).

38. Mostowy, R. J. & Holt, K. E. Diversity-generating machines: genetics of bacterial sugar-coating. Trends Microbiol. xx, 1–14 (2018).

39. Lin, T. L. et al. Isolation of a bacteriophage and its depolymerase specific for K1 capsule of *Klebsiella pneumoniae*: implication in typing and treatment. J Infect Dis. 210, 1734–1744 (2014).

40. Rieger-Hug, D. & Stirm, S. Comparative study of host capsule depolymerases associated with *Klebsiella* bacteriophages. Virology 113, 363–378 (1981).

41. Rodriguez-Valera, F. et al. Explaining microbial population genomics through phage predation. Nat Rev Microbiol. 7, 828–836 (2009).

42. Wyres, K. L. & Holt, K. E. *Klebsiella pneumoniae* as a key trafficker of drug resistance genes from environmental to clinically important bacteria. Curr. Opin. Microbiol. 45, 131–139 (2018).

43. Bidewell, C. A. et al. Emergence of *Klebsiella pneumoniae* subspecies *pneumoniae* as a cause of septicaemia in pigs in England. PLoS One 13, e0191958 (2018).

44. Comandatore, F. et al. Gene composition as a potential barrier to large recombinations in the bacterial pathogen *Klebsiella pneumoniae*. bioRxiv (2018).

45. Page, A. J. et al. Roary: rapid large-scale prokaryote pan genome analysis. Bioinformatics btv421 (2015). doi:10.1093/bioinformatics/btv421

46. Tettelin, H., Riley, D., Cattuto, C. & Medini, D. Comparative genomics: the bacterial pan-genome. Curr Op Microbiol. 11, 472–477 (2008).

47. Roux, S., Enault, F., Hurwitz, B. L. & Sullivan, M. B. VirSorter: mining viral signal from microbial genomic data. PeerJ 3, e985 (2015).

48. Arredondo-Alonso, S., Willems, R. J., van Schaik, W. & Schürch, A. C. On the (im)possibility of reconstructing plasmids from whole-genome short-read sequencing data. Microb. Genomics 3, (2017).

49. Carattoli, A. et al. PlasmidFinder and pMLST: *in silico* detection and typing of plasmids. Antimicrob Agents Chemother. 58, 3895–3903 (2014).

50. Orlek, A. et al. Ordering the mob: Insights into replicon and MOB typing schemes from analysis of a curated dataset of publicly available plasmids. Plasmid 91, 42–52 (2017).

51. Francia, M. V. et al. A classification scheme for mobilization regions of bacterial plasmids. FEMS Microbiol. Rev. 28, 79–100 (2004).

52. Chen, L., Mathema, B., Pitout, J. D., DeLeo, F. R. & Kreiswirth, B. N. Epidemic *Klebsiella pneumoniae* ST258 Is a hybrid strain. MBio 5, e01355–14 (2014).

53. Bowers, J. R. et al. Genomic analysis of the emergence and rapid global dissemination of the clonal group 258 *Klebsiella pneumoniae* pandemic. PLoS One 10, e0133727 (2015).

54. Martin, J. et al. Covert dissemination of carbapenemase-producing *Klebsiella pneumoniae* (KPC) in a successfully controlled outbreak: long- and short-read whole-genome sequencing demonstrate multiple genetic modes of transmission. J Antimicrob Chemother. 72, 3025–3034 (2017).

55. Sheppard, A. E. et al. Nested Russian doll-like genetic mobility drives rapid dissemination of the carbapenem resistance gene *bla*KPC. Antimicrob Agents Chemother. 60, 3767–3778 (2016).

56. Cheng, H. Y. et al. RmpA regulation of capsular polysaccharide biosynthesis in *Klebsiella pneumoniae* CG43. J Bacteriol. 192, 3144–3158 (2010).

57. Wei, D., Yuminaga, Y., Shi, J. & Hao, J. Non-capsulated mutants of a chemical-producing *Klebsiella pneumoniae* strain. Biotechnol Lett 40, 679–687 (2018).

58. Chewapreecha, C. et al. Dense genomic sampling identifies highways of pneumococcal recombination. Nat Genet. 46, 305–309 (2014).

59. Clements, A. et al. The major surface-associated saccharides of <I>Klebsiella pneumoniae</i> contribute to host cell association. PLoS One 3, e3817 (2008).

60. Anthony, K. G., Sherburne, C., Sherburne, R. & Frost, L. S. The role of the pilus in recipient cell recognition during bacterial conjugation mediated by F-like plasmids. Mol. Microbiol. 13, 939–953 (1994).

61. Pérez-Mendoza, D. & de la Cruz, F. *Escherichia coli* genes affecting recipient ability in plasmid conjugation: Are there any? BMC Genomics 10, 71 (2009).

62. Pan, Y.-J. et al. Genetic analysis of capsular polysaccharide synthesis gene clusters in 79 capsular types of *Klebsiella* spp. Nat Sci Rep. 5, 15573 (2015).

63. Fournet-Fayard, S., Joly, B. & Forestier, C. Transformation of wild type *Klebsiella pneumoniae* with plasmid DNA by electroporation. J. Microbiol. Methods 24, 49–54 (1995).

64. Conlan, S. et al. Plasmid dynamics in KPC-positive *Klebsiella pneumoniae* during long-term patient colonization. 7, e00742–16 (2016).

65. Löhr, I. H. et al. Persistence of a pKPN3-like CTX-M-15-encoding IncFIIK plasmid in a *Klebsiella pneumoniae* ST17 host during two years of intestinal colonization. PLoS One 10, e0116516 (2015).

66. Buckner, M. M. C. et al. Clinically relevant plasmid-host interactions indicate that transcriptional and not genomic modifications ameliorate fitness costs of *Klebsiella pneumoniae* carbapenemase-carrying plasmids. MBio 9, e02303–17 (2018).

67. Du, P., Zhang, Y. & Chen, C. Emergence of carbapenem-resistant hypervirulent Klebsiella pneumoniae. Lancet Infect. Dis. 3099, 30629 (2017).

68. Lam, M. C. C., Wick, R. R., Wyres, K. L. & Holt, K. E. Kleborate: comprehensive genotyping of *Klebsiella pneumoniae* genome assemblies. (2018). at <https://github.com/katholt/Kleborate>

69. Gorrie, C. L. et al. Antimicrobial resistant *Klebsiella pneumoniae* carriage and infection in specialized geriatric care wards linked to acquisition in the referring hospital. Clin. Infect. Dis. (2018). doi:10.1093/cid/ciy027

70. The, H. C. et al. A high-resolution genomic analysis of multidrug-resistant hospital outbreaks of *Klebsiella pneumoniae*. EMBO Molec Med. 7, 227–239 (2015).

71. Struve, C. et al. Mapping the evolution of hypervirulent *Klebsiella pneumoniae*. MBio. 6, 1–12 (2015).

72. Stoesser, N. et al. Predicting antimicrobial susceptibilities for *Escherichia coli* and *Klebsiella pneumoniae* isolates using whole genomic sequence data. J Antimicrob Chemother. 68, 2234–2244 (2013).

73. Stoesser, N. et al. Genome sequencing of an extended series of NDM-producing *Klebsiella pneumoniae* isolates from neonatal infections in a Nepali hospital characterizes the extent of community-versus hospital-associated transmission in an endemic setting. Antimicrob Agents Chemother. 58, 7347–7357 (2014).

74. Wand, M. E. et al. Characterization of pre-antibiotic era *Klebsiella pneumoniae* Isolates with respect to antibiotic/disinfectant susceptibility and virulence in *Galleria mellonella*. Antimicrob Agents Chemother. 59, 3966–3972 (2015).

75. Deleo, F. R. et al. Molecular dissection of the evolution of carbapenem-resistant multilocus sequence type 258 *Klebsiella pneumoniae*. Proc Natl Acad Sci U S A. 111, 4988–4993 (2014).

76. Davis, G. S. et al. Intermingled *Klebsiella pneumoniae* populations between retail meats and human urinary tract infections. Clin Infect Dis 61, 892–899 (2015).

77. Langmead, B. & Salzberg, S. L. Fast gapped-read alignment with Bowtie 2. Nat Methods. 9, 357–359 (2012).

78. Li, H. et al. The Sequence Alignment/Map format and SAMtools. Bioinformatics 25, 2078–2079 (2009).

79. Price, M. N., Dehal, P. S. & Arkin, A. P. FastTree: computing large minimum evolution trees with profiles instead of a distance matrix. Mol Biol Evol. 26, 1641–1650 (2009).

80. Assefa, S., Keane, T. M., Otto, T. D., Newbold, C. & Berriman, M. ABACAS: algorithm-based automatic contiguation of assembled sequences. Bioinformatics 25, 1968–1969 (2009).

81. Carver, T. J. et al. ACT: the Artemis comparison tool. Bioinformatics 21, 3422–3423 (2005).

82. Gorrie, C. L. et al. Gastrointestinal carriage is a major reservoir of *K. pneumoniae* infection in intensive care patients. Clin Infect Dis. Epub ahead of print (2017). doi:10.1101/096446

83. Wick, R. R., Judd, L. M., Gorrie, C. L. & Holt, K. E. Completing bacterial genome assemblies with multiplex MinION sequencing. MGen. 3, (2017).

84. Wick, R. R., Judd, L. M., Gorrie, C. L. & Holt, K. E. Unicycler: resolving bacterial genome assemblies from short and long sequencing reads. PLoS Comp Biol. 13, e1005595 (2017).

85. Inouye, M. et al. SRST2: Rapid genomic surveillance for public health and hospital microbiology labs. Genome Med. 6, 90 (2014).

86. Bankevich, A. et al. SPAdes: a new genome assembly algorithm and its applications to single-cell sequencing. J Comput Biol. 19, 455–477 (2012).

87. Wick, R. R., Heinz, E., Holt, K. E. & Wyres, K. L. Kaptive Web: user-friendly capsule and lipopolysaccharide serotype prediction for *Klebsiella* genomes. J Clin Microbiol. 56, e00197–18 (2018).

88. Oksanen, J. et al. vegan: community ecology package. R package v 2.4.3. (2017).

89. Jost, L. Entropy and diversity. Oikos 113, 363–375 (2006).

90. Stamatakis, A. RAxML-VI-HPC: maximum likelihood-based phylogenetic analyses with thousands of taxa and mixed models. Bioinformatics. 22, 2688–2690 (2006).

91. Seemann, T. Prokka: rapid prokaryotic genome annotation. Bioinformatics 30, 2068–2069 (2014).

92. Jombart, T. Adegenet: A R package for the multivariate analysis of genetic markers. Bioinformatics 24, 1403–1405 (2008).

93. Snipen, L. & Liland, K. H. micropan: An R-package for microbial pan-genomics. BMC Bioinformatics 16, 79 (2015).

94. Wood, D. E. & Salzberg, S. L. Kraken: Ultrafast metagenomic sequence classification using exact alignments. Genome Biol. 15, R46 (2014).

95. Fu, L., Niu, B., Zhu, Z., Wu, S. & Li, W. CD-HIT: Accelerated for clustering the next-generation sequencing data. Bioinformatics 28, 3150–3152 (2012).

96. Bland, C. et al. CRISPR recognition tool (CRT): a tool for automatic detection of clustered regularly interspaced palindromic repeats. BMC Bioinformatics 8, 209 (2007).

97. Ostria-Hernández, M. L., Sánchez-Vallejo, C. J., Ibarra, J. A. & Castro-Escarpulli, G. Survey of clustered regularly interspaced short palindromic repeats and their associated Cas proteins (CRISPR/Cas) systems in multiple sequenced strains of *Klebsiella pneumoniae*. BMC Res Notes 8, 332 (2015).

98. Burstein, D. et al. New CRISPR–Cas systems from uncultivated microbes. Nature 542, 237–241 (2017).

99. Eddy, S. R., Wheeler, T. J. & HMMER development team. HMMER3 Suite. (2015). at <http://hmmer.org/>

100. Shen, J., Lv, L., Wang, X., Xiu, Z. & Chen, G. Comparative analysis of CRISPR-Cas systems in *Klebsiella* genomes. J. Basic Microbiol. 1–12 (2017). doi:10.1002/jobm.201600589

101. Oliveira, P. H., Touchon, M. & Rocha, E. P. C. Regulation of genetic flux between bacteria by restriction–modification systems. Proc Natl Acad Sci U S A. 113, 5658–5663 (2016).

102. Cury, J., Jové, T., Touchon, M., Néron, B. & Rocha, E. P. Identification and analysis of integrons and cassette arrays in bacterial genomes. Nucleic Acids Res. 44, 4539–4550 (2016).

103. Henson, S. P. et al. Molecular epidemiology of *Klebsiella pneumoniae* invasive infections over a decade at Kilifi County Hospital in Kenya. Int J Med Microbiol. 307, 422–429 (2017).

104. Runcharoen, C. et al. Whole genome sequencing reveals high-resolution epidemiological links between clinical and environmental *Klebsiella pneumoniae*. Genome Med 9, 6 (2017).

105. Moradigaravand, D., Martin, V., Peacock, S. J. & Parkhill, J. Evolution and epidemiology of multidrug-resistant *Klebsiella pneumoniae* in the United Kingdom. MBio 8, e01976–16 (2017).

106. Smit, P. et al. Transmission patterns of hyper-endemic multi-drug resistant *Klebsiella pneumoniae* in a Cambodian neonatal unit: a longitudinal study with whole genome sequencing. bioRxiv (2017).

107. Bouckaert, R. et al. BEAST 2: A software platform for Bayesian evolutionary analysis. PLoS Comp Biol. 10, e1003537 (2014).

